# Mitochondrial network branching enables rapid protein spread with slower mitochondrial dynamics

**DOI:** 10.1101/2024.05.07.593000

**Authors:** Prabha Chuphal, Aidan I. Brown

## Abstract

Mitochondrial network structure is controlled by the dynamical processes of fusion and fission, which merge and split mitochondrial tubes into structures including branches and loops. To investigate the impact of mitochondrial network dynamics and structure on the spread of proteins and other molecules through mitochondrial networks, we used stochastic simulations of two distinct quantitative models that each included mitochondrial fusion and fission, and particle diffusion via the network. Better-connected mitochondrial networks and networks with faster dynamics exhibit more rapid particle spread on the network, with little further improvement once a network has become well-connected. As fragmented networks gradually become better-connected, particle spread either steadily improves until the networks become well-connected for slow-diffusing particles or plateaus for fast-diffusing particles. We compared model mitochondrial networks with both end-to-end and end-to-side fusion, which form branches, to non-branching model networks that lack end-to-side fusion. To achieve the optimum (most rapid) spread that occurs on well-connected branching networks, non-branching networks require much faster fusion and fission dynamics. Thus the process of end-to-side fusion, which creates branches in mitochondrial networks, enables rapid spread of particles on the network with relatively slow fusion and fission dynamics. This modeling of protein spread on mitochondrial networks builds towards mechanistic understanding of how mitochondrial structure and dynamics regulate mitochondrial function.

## I INTRODUCTION

Mitochondria are highly dynamic organelles with multiple important functions [1, 2], notably production of ATP via oxidative phosphorylation [3]. Mitochondrial dysfunction is connected to several diseases, such as diabetes, cancer, and Parkinson’s disease, among others [4– 7]. While often depicted as bean-shaped puncta, mitochondrial fusion and fission dynamics allow formation of mitochondrial tubes of a range of sizes [8]. Tip-to-side mitochondrial fusion forms three-way junctions that branch mitochondrial tubes, allowing mitochondria to form extended network structures that include loops [9– 11]. Mitochondrial dynamics are important to mitochondrial quality control and function [12], as well as to cell metabolism [13], motion [14], and differentiation [15].

The level of connectivity for mitochondria is controlled by both fission and fusion events [16–19], as well as organelle movement [20–22]. Relatively frequent fission leads to more fragmented mitochondria, while relatively frequent fusion leads to a more connected mitochondrial network. Mitochondria in yeast cells can be well-connected, with a mean of two mitochondrial fragments and most mitochondrial mass part of a relatively large mitochondrial fragment [9]; mitochondrial fragments in mammalian cells can take a range of sizes from large to small [8]; while in plant cells mitochondria are typically many small fragments [23]. Mitochondrial network connectivity can be adjusted by changes to metabolism and cell stress [24–30]. Various hypotheses have been explored to explain why mitochondria form extended and dynamic connected networks [31, 32], including to share proteins and other molecules [33], undergoing fission to provide growing and dividing cells with sufficient numbers of mitochondria [34], and to help separate poorly-functioning mitochondria to be removed by the mitophagy quality control pathway [1, 12].

We will focus on how mitochondrial network connectivity and dynamics facilitate the sharing of proteins and other molecules, which appears key to providing mitochondria with a full protein complement. Most mitochondrial proteins are encoded by nuclear DNA, and must be imported into mitochondria [35]. Low mRNA copy number for some genes combined with co-translational delivery and stochastic targeting to mitochondria can lead some mitochondrial fragments to directly receive few copies of a specific protein — these proteins could instead be provided following fusion with another fragment [36, 37]. A small number of key mitochondrial genes are encoded by mitochondrial DNA, of which there are many copies in each cell [38]. With limited mitochondrial DNA error-correction mechanisms, mutation levels are relatively high [38]. While proteins produced in one mitochondrial fragment may be mutated and dysfunctional, functional proteins produced in another fragment can be shared following fusion, enabling local function, known as complementation [39]. While fusing all mitochondria together into one network may facilitate rapid sharing, this could hamper the distribution of mitochondria throughout the cell and may pose challenges for other purposes served by mitochondrial dynamics [32].

Recently-developed experimental techniques enable measurement of mitochondrial network structure and dynamics [40]. Additionally, experiments can measure localization of mRNA for nuclear-encoded mitochondrial proteins [29, 41], mitochondrial nucleoid locations [42], and mitochondrial protein concentrations [37]. Beyond directly advancing understanding of many aspects of mitochondria, these measurements provide a quantitative basis to model mitochondrial networks and processes, including those important to sharing proteins among mitochondria. Understanding how mitochondrial dynamics control network structure and connectivity changes is important to describing protein spread in mitochondria. Modeling work has outlined how a balance between fusion and fission leads to critical mitochondrial networks near the percolation threshold [8, 43, 44], how mitochondrial motion along filaments is key to spatially organizing mitochondrial networks [45], and very recent work explored how kinetics and mechanics control spatial mitochondrial network morphology and found high dependence of morphology and rearrangement timescales on fission and fusion rates [19].

To address how mitochondrial structure affects protein spread on a static network, recent modeling work showed that mitochondrial network snapshots that contain loops allow faster diffusive search compared to networks without loops [9, 10]. For spread on dynamic mitochondrial networks, a two-dimensional lattice model showed stronger dependence of effective diffusivity on connectivity for slower dynamics [31], a two-dimensional spatial model with mitochondrial motion along filaments suggested the rate of mitochondrial fusion impacts mitochondrial health [21], an aspatial model indicated more frequent fusion suppresses protein concentration fluctuations between mitochondria [37], and a model of heavily-fragmented mitochondria suggested mitochondrial DNA gene products efficiently spread (using a contact measure of spread) for parameters describing living plant cells [46]. These studies of dynamic networks considered spread through the mitochondrial network in a specific limit or as a secondary focus, and we are unaware of other work that focuses on how mitochondrial dynamics affect protein spread through the mitochondrial networks. In particular, there appears to be little to no work on how spread is impacted by the tip-to-side fusion processes that form branches in mitochondrial networks.

To investigate how mitochondrial network structure and dynamics, particularly branching of networks, affect protein spread, we simulate two different stochastic models of mitochondrial networks, one on a two-dimensional spatial lattice and a second agent-based aspatial model. Among many similarities, the particle spread behavior of both models suggests that the branching of mitochondrial networks allows for rapid spread with much slower mitochondrial dynamics compared to networks without branching.

## II RESULTS

### A Two-dimensional lattice-based model

We first explore a mitochondrial network model with stochastic fusion and fission between mitochondrial units embedded on a two-dimensional lattice (see Fig. 1, left panel), following earlier work [31]. Although embedded in a three-dimensional volume, the spatial arrangement of mitochondria (in flat cells [21] or in a layer just inside the yeast cell membrane [9]) and analysis of mitochondrial networks (percolation exponents closer to those expected for two dimensions than three dimensions [44]) suggests consideration of two-dimensional mitochondrial networks.

**FIG. 1.**
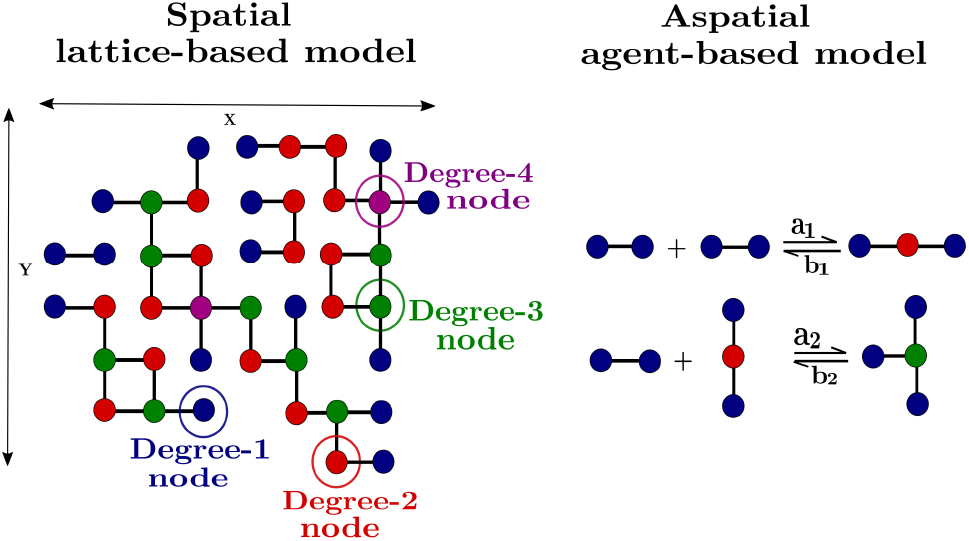
Schematic representing two mitochondrial network models. Left: Spatial lattice-based model with mitochondria represented by nodes on a two-dimensional lattice. Neighboring nodes connect (fuse) at rate *λ*_add_ and disconnect (fission) at rate *λ*_remove_. Right: Aspatial agent-based model with each minimal mitochondrial fragment represented by two permanently-connected nodes. Two free mitochondrial ends (only joined to permanently-connected partner) fuse into a degree-two node with rate *a*_1_ and a degree-two node fissions into two free ends with rate *b*_1_; a free end and degree-two node fuse into a degree-three node with rate *a*_2_ and degree three nodes fission into a free end and a degree-two node with rate *b*_2_. For both panels, node color indicates node degree.

Neighboring nodes connect (fuse) at rate *λ*_add_ and disconnect (fission) at rate *λ*_remove_, such that neighboring mitochondria are connected with probability *p* = *λ*_add_*/*(*λ*_add_ + *λ*_remove_). On a fully-connected network, a particle can move on the network with effective diffusivity *D*_eff_ equal to the particle diffusivity *D* = 1, such that the rate for moving to each neighboring node is one, setting the timescale. For an incomplete network, the particle would be unable to step across missing connections, so *D*_eff_ *< D*. Further lattice simulation details are described in the Appendix.

Figure 2A shows that the effective diffusivity increases as the connection probability *p* increases, recapitulating earlier work [31]. For small *p* the network mostly contains small node islands or isolated nodes, while for large *p* most nodes are connected into a single component (Fig. 2A insets). With *τ* = 1*/*(*λ*_add_ + *λ*_remove_) the timescale of network dynamics, fast dynamics (small *τ*) leads the effective diffusivity to increase close to linearly with increasing connection probability *p*, with faster dynamics more linear with a higher *D*_eff_ at a given *p*. For infinitely fast dynamics *τ* → 0 each attempted step by the particle occurs with probability *p*, uncorrelated with previous steps. For slow dynamics (large *τ*) the effective diffusivity increases very little below the two-dimensional percolation threshold [47] *p* = 0.5 and rapidly rises above *p* = 0.5. This reflects the longer lifetime of connections and disconnections, such that successful and unsuccessful attempted steps become increasingly correlated in time. For *τ* → ∞ the network is frozen, with no changes in connections. For *τ* → ∞, Fig. 2A shows the effective diffusivity does not quite reach zero at and immediately below *p* = 0.5 due to the finite system simulated, while the effective diffusivity would sharply depart from zero at this threshold on an infinite network [47].

**FIG. 2.**
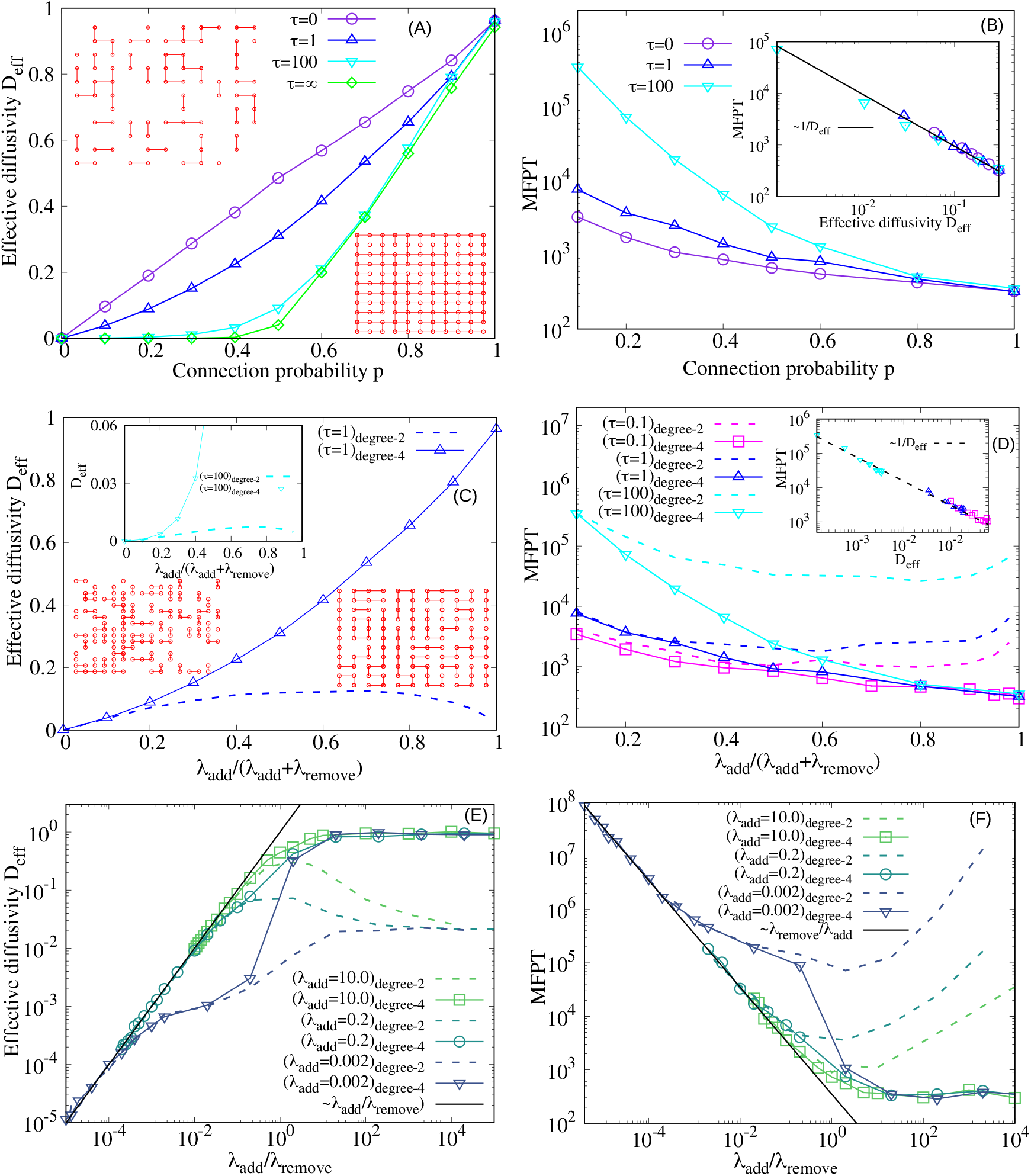
Effective diffusivity and search time for particle diffusing on the model dynamic mitochondrial network on a two-dimensional lattice. (A) The effective diffusivity *D*_eff_ as connection probability *p* is varied. Legend indicates network dynamics timescale *τ*. Insets show typical networks for *p* = 0.2 and *p* = 0.9. (B) The mean first-passage time (MFPT) of the diffusing particle to find randomly-selected lattice site from the initial position at another randomly-selected lattice site, as connection probability *p* is varied. Inset shows MFPT vs *D*_eff_ from main plots of panels A and B. Main plot legend, indicating *τ*, also applies to inset, with inset black line indicating MFPT ∼ 1*/D*_eff_ trend. In panels, A and B, nodes are permitted to connect to all four nearest neighbors. (C) *D*_eff_ as fusion fraction of dynamics *λ*_add_*/*(*λ*_add_ + *λ*_remove_) is varied. The main plot is for *τ* = 1 and inset is for *τ* = 100. (D) MFPT of the diffusing particle to find randomly-selected lattice site from the initial position at another randomly-selected lattice site, as *λ*_add_*/*(*λ*_add_ + *λ*_remove_) is varied. Legend indicates *τ*. Inset shows degree-2 MFPT vs *D*_eff_ from main plots of panels C and D. Main plot legend also applies to inset, with inset black curve indicating MFPT ∼ 1*/D*_eff_ trend. (E) *D*_eff_ as balance between fusion and fission *λ*_add_*/λ*_remove_ is varied. Legend indicates fusion rate *λ*_add_ and black curve is ∼ *λ*_add_*/λ*_remove_ trend. (F) MFPT as *λ*_add_*/λ*_remove_ is varied. Legend indicates *λ*_add_ and black curve is ∼ *λ*_remove_*/λ*_add_ trend. In panels C – F, solid curves are for networks that permit nodes to connect to all four nearest neighbors (degree-4), and dashed curves indicate networks that permit nodes to connect to a maximum of two nearest neighbors (degree-2), as indicated in legends. Across all panels, particle diffusivity *D* = 1, the lattice is 10 by 10 with nodes on a square grid with intervals of one, and data averaged over 100 samples.

**FIG. 3.**
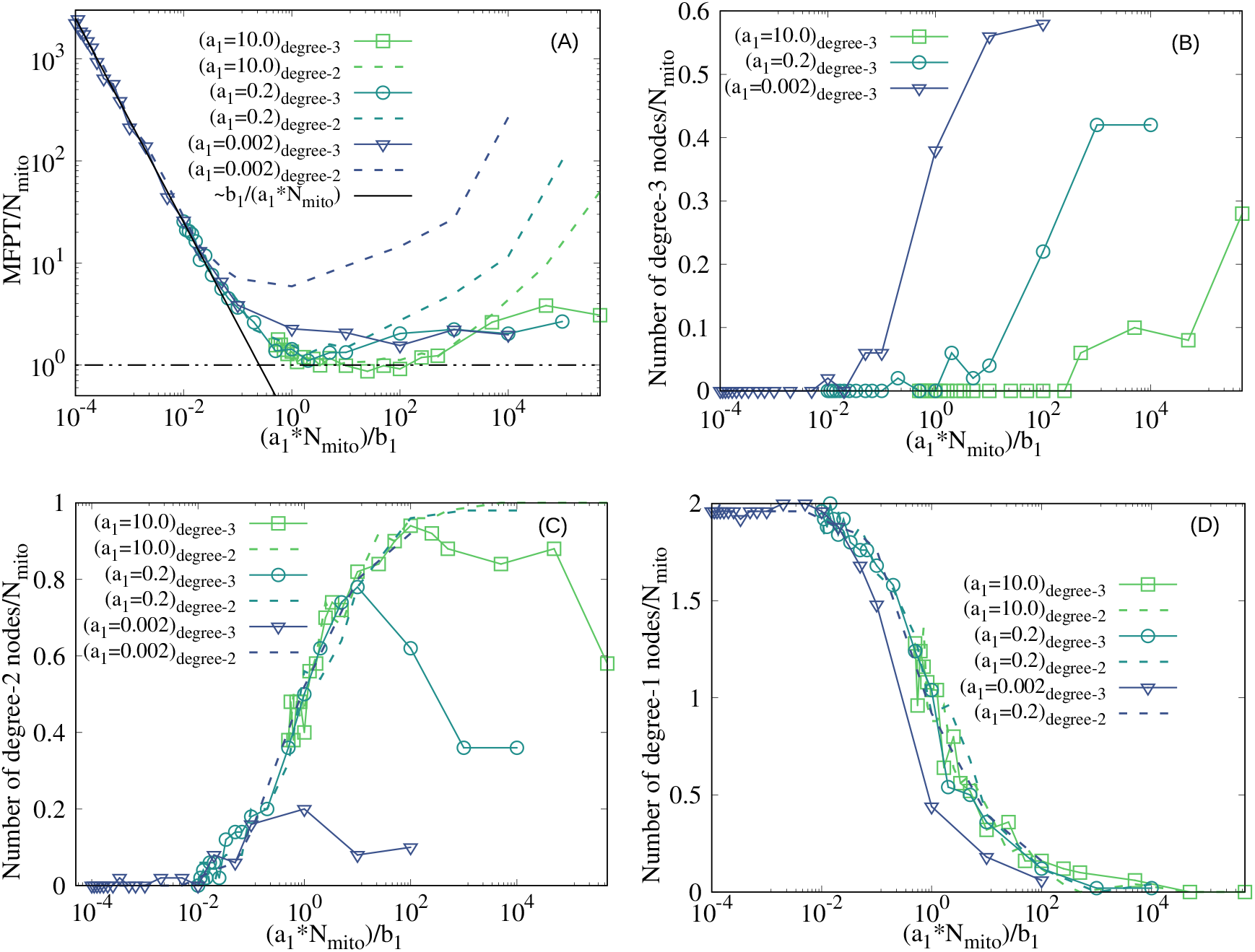
Search times for particle diffusing on and network structure of the model aspatial dynamic mitochondrial network. (A) Mean first-passage time (MFPT) of a particle to find a randomly-selected mitochondrial fragment from an initial location on another randomly-selected fragment, as the scaled ratio of end-to-end fusion to fission *a*_1_*N*_mito_*/b*_1_ is varied. Solid black line is MFPT ∼ *b*_1_*/*(*a*_1_*N*_mito_) trend and dot-dashed black line is minimum search time ⟨*T*_min_⟩ from Eq. 4. For all panels, solid curves are for branching (degree-3, end-to-side fusion rate *a*_2_ = 0.01) networks, and dashed curves are non-branching (degree-2, *a*_2_ = 0) networks, and legend further indicates end-to-end fusion rate *a*_1_. (B,C,D) The scaled number of (B) three-way junctions (degree-three nodes), (C) end-to-end fusions (degree-two nodes, solid lines with branching and dashed lines without branching), and (D) free ends (degree-one nodes, solid lines with branching and dashed lines without branching) as *a*_1_*N*_mito_*/b*_1_ is varied, with network branching via end-to-side fusion. Across all panels, the end-to-side fusion rate for branching (degree-3) networks is *a*_2_ = 0.01, the rate to move to connected mitochondrial fragment is *k*_m_ = 1, the number of minimal mitochondrial fragments *N*_mito_ = 50, and data averaged over 100 samples.

Complementing the effective diffusivity of Fig. 2A, Fig. 2B shows the time to first arrive at a target node in the network, the mean first-passage time (MFPT), with randomly selected initial and target nodes. The MFPT decreases with increasing connection probability *p*, with shorter MFPT for faster dynamics (smaller *τ*). The MFPT for large *τ* decreases by multiple orders of magnitude as *p* increases from 0.1 to 1 because for low *p* the particle must wait for fusion events to enable movement to new nodes, and *τ* represents this waiting time for a given pair of neighboring mitochondria. In contrast, the MFPT for small *τ* (including the infinitely-fast dynamics of *τ* = 0) decreases by approximately an order of magnitude from *p* = 0.1 to *p* = 1 as the search is slowed by low *p* due to less connectivity, rather than long waiting times for dynamics. As the effective diffusivity is essentially the probability for the particle to successfully step from one mitochondria to another, allowing the particle to proceed on a search, the MFPT inversely scales with the effective diffusivity (Fig. 2B inset).

The results of Fig. 2A,B permit mitochondrial nodes on the two-dimensional lattice to connect to all four of their nearest neighbors, allowing degree-four nodes. We now examine how restricting the maximum node degree to below four impacts diffusion and search. On networks with such a restriction, the connection probability *p*≠ *λ*_add_*/*(*λ*_add_ + *λ*_remove_), as a fourth connection to a particular node cannot form. For networks with a maximum node degree of three (approximately as observed in mitochondrial networks [9]), the effective diffusivity *D*_eff_ and MFPT are qualitatively similar to networks permitting degree-four nodes, with monotonic changes in both quantities (see Fig. 5). Restricting to a maximum node degree of three lowers the effective diffusivity and raises the MFPT compared to degree-four networks for better-connected networks with larger *λ*_add_*/*(*λ*_add_ + *λ*_remove_).

In contrast, networks with a maximum node degree of two exhibit qualitative distinctions in effective diffusivity and search behavior compared to networks permitted degree-three and degree-four nodes (Fig. 2C,D). Such networks with a maximum node degree of two represent hypothetical mitochondrial networks without tip-to-side fusion or the associated network branching. For low *λ*_add_*/*(*λ*_add_ + *λ*_remove_) ≲ 0.2 the effective diffusivity *D*_eff_ for networks permitting maximum degree-two nodes is similar to the *D*_eff_ with degree-four nodes (Fig. 2C). For *λ*_add_*/*(*λ*_add_ + *λ*_remove_) ≳ 0.2 the *D*_eff_ for maximum degree-two networks increases much more slowly as *λ*_add_*/*(*λ*_add_ + *λ*_remove_) increases than for degree-four networks. Although *D*_eff_ for degree-four networks monotonically increases with *λ*_add_*/*(*λ*_add_ + *λ*_remove_), for high *λ*_add_*/*(*λ*_add_+*λ*_remove_) ≳ 0.7 the *D*_eff_ for maximum degree-two networks decreases with higher *λ*_add_*/*(*λ*_add_+*λ*_remove_). Larger *τ* = 1*/*(*λ*_add_ + *λ*_remove_), representing slower network rearrangement, leads to greater difference between degree-four and degree-two *D*_eff_ (Fig. 2C main plot compared with inset plot).

Consistent with the trend for effective diffusivity, Fig. 2D shows similar and decreasing MFPT for degree-four and degree-two networks for small and increasing *λ*_add_*/*(*λ*_add_ + *λ*_remove_), higher MFPT for degree-two than degree-four networks at intermediate *λ*_add_*/*(*λ*_add_ + *λ*_remove_), and increasing MFPT for degree-two networks for high and increasing *λ*_add_*/*(*λ*_add_ + *λ*_remove_) values. The difference between degree-four and degree-two MFPT also increased with larger *τ*. The inset of Fig. 2D shows a 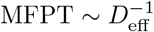 relationship for degree-two networks consistent with the relationship found in the inset of Fig. 2B for degree-four networks. Similar results to Fig. 2A,C are found for a larger lattice (see Fig. 4 in Appendix).

**FIG. 4.**
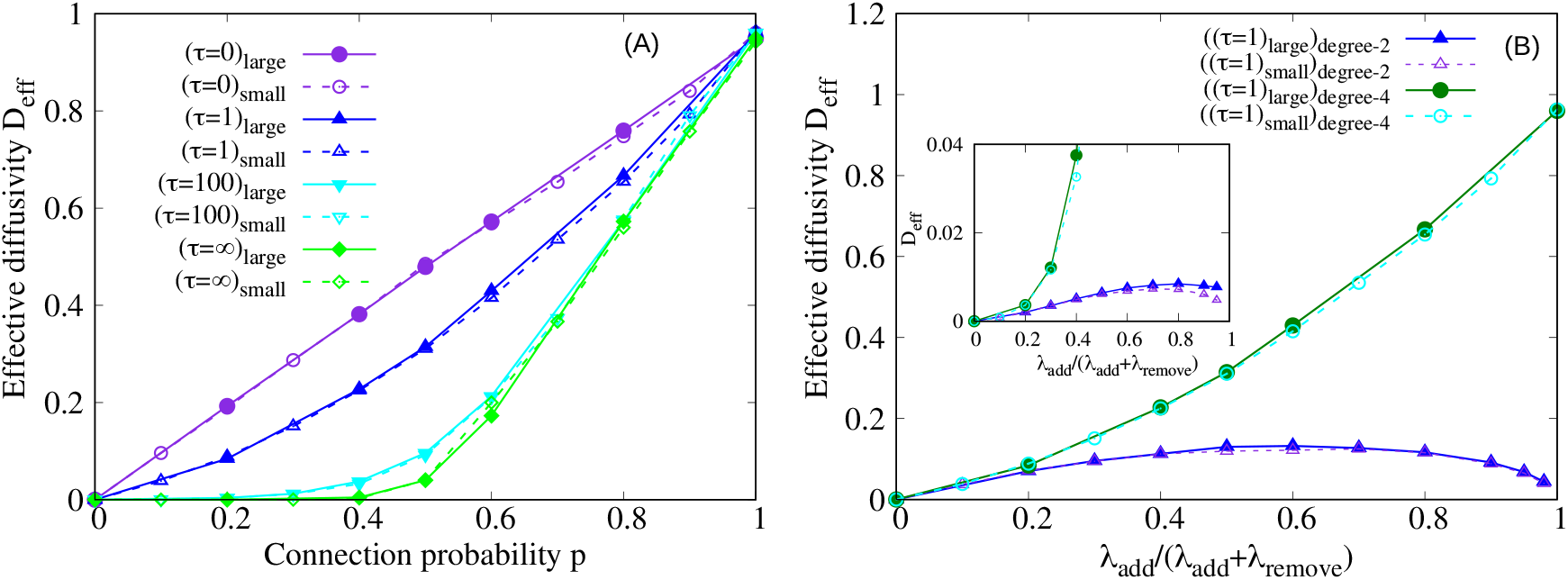
Comparison of effective diffusivity *D*_eff_ for two different system sizes. (A) Effective diffusivity *D*_eff_ of a particle diffusing on the model dynamic mitochondrial network on a two-dimensional lattice as connection probability *p* is varied. Solid curves with filled markers are for a 20 by 20 lattice and dashed curves with open markers are for a 10 by 10 lattice, with nodes on a square grid with intervals of one. Legend indicates network dynamics timescale *τ*. Particle diffusivity *D* = 1 and data averaged over 100 samples. (B) Effective diffusivity *D*_eff_ for particle diffusing on model dynamic mitochondrial network on two-dimensional lattice as fusion fraction of dynamics *λ*_add_*/*(*λ*_add_ + *λ*_remove_) is varied, for 10 by 10 (small, dashed curves) and 20 by 20 (large, solid curves) systems. System size and whether nodes can connect to all four nearest neighbors (degree-4) or only two nearest neighbors (degree-2). The main plot is for network dynamics timescale *τ* = 1 and the inset is for *τ* = 100. Legend applies to both the main plot and the inset. Data averaged over 100 samples.

**FIG. 5.**
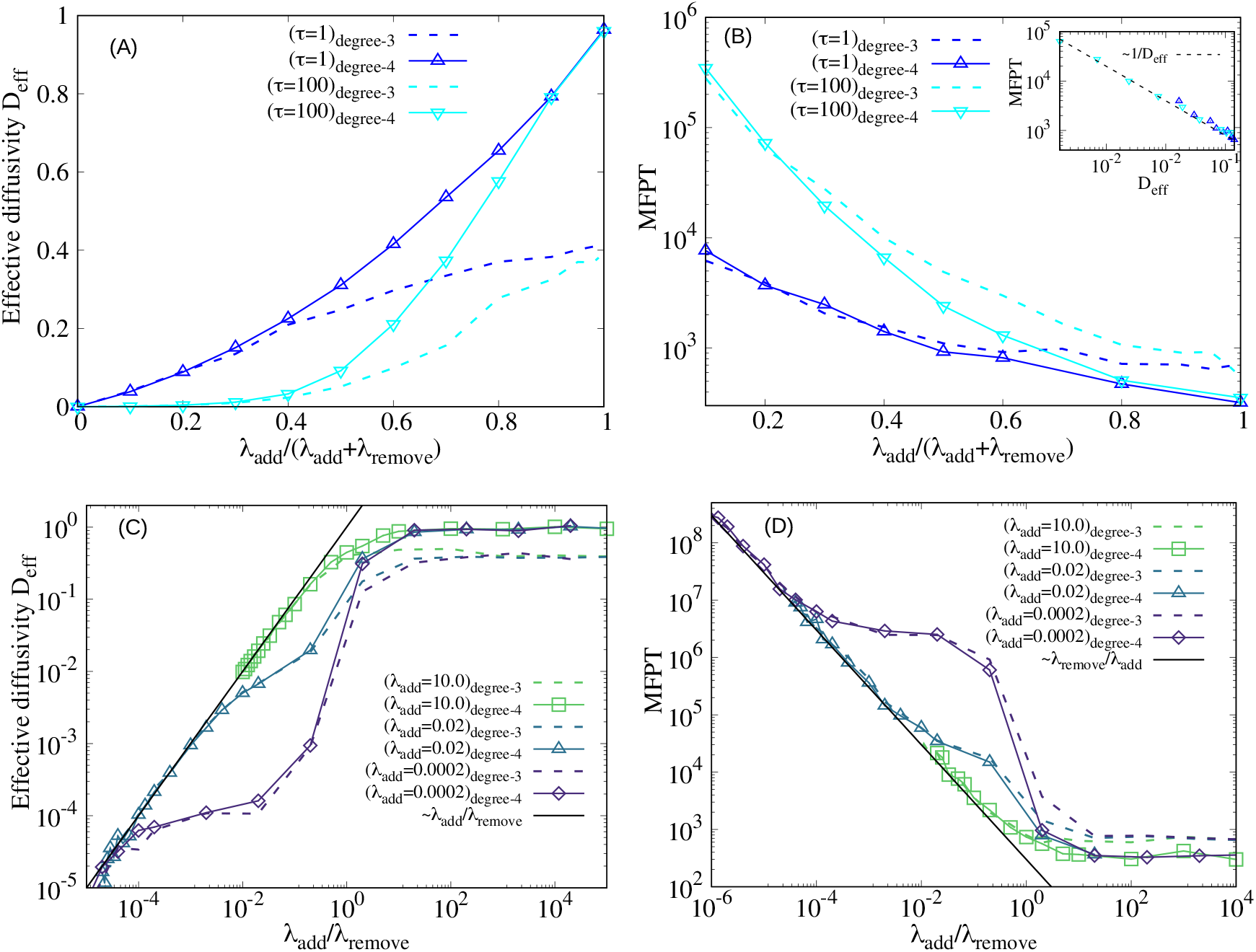
Comparison of particle diffusivity and search time on lattice networks permitting degree-four and degree-three nodes. (A) Effective diffusivity *D*_eff_ as fusion fraction of dynamics *λ*_add_*/*(*λ*_add_ + *λ*_remove_) is varied. Across panels, the legend indicates network dynamics timescale *τ* and whether nodes can connect to all four nearest neighbors (degree-4, solid curves) or only three nearest neighbors (degree-3, dashed curves). (B) Mean first passage time (MFPT) as *λ*_add_*/*(*λ*_add_ + *λ*_remove_) is varied. Inset shows MFPT vs *D*_eff_ from main plots of panels A and B. Dashed black line in the inset shows ∼ 1*/D*_eff_ trend. (C) Effective diffusivity *D*_eff_ as balance between fusion and fission *λ*_add_*/λ*_remove_ is varied. Black line is ∼ *λ*_add_*/λ*_remove_ trend. (D) MFPT as *λ*_add_*/λ*_remove_ is varied. Black line is ∼ *λ*_remove_*/λ*_add_ trend. Across all panels, particle diffusivity *D* = 1, the lattice is 10 by 10 with nodes on a square grid with intervals of one, and data averaged over 100 samples.

Particles have similar effective diffusivity and MFPT on networks with degree-four or maximum of degree-two nodes for small *λ*_add_*/*(*λ*_add_ + *λ*_remove_) because in this low-connectivity regime limiting node degree to a maximum of two affects few nodes as they are already largely disconnected (see left network insets of Fig. 2A,C). As *λ*_add_*/*(*λ*_add_ + *λ*_remove_) increases beyond this low-connectivity regime, node connections are decreased by limiting node degree to a maximum of two, reducing effective diffusivity and increasing MFPT in these networks compared to networks without this constraint. As *λ*_add_*/*(*λ*_add_ + *λ*_remove_) exceeds approximately 0.7, further increase in *λ*_add_*/*(*λ*_add_ + *λ*_remove_) decreases effective diffusivity and increases MFPT if node degree is restricted to a maximum of two because the network freezes into linear or looped fragments (see right network inset of Fig. 2C). Nodes that already have two connections to neighboring nodes cannot add a third connection, causing the formation of new connections to be coupled to the preceding removal of existing connections. At a fixed timescale of network dynamics *τ* = 1*/*(*λ*_add_ + *λ*_remove_), increasing *λ*_add_*/*(*λ*_add_ + *λ*_remove_) reduces *λ*_remove_, slowing the rearrangement of the network. Particle confinement to a particular linear or looped fragment until the local network rearranges manifests as decreased diffusivity and increased MFPT compared to a less connected network with more rapid rearrangement dynamics. There is a smaller difference between degree-four and maximum degree-two networks for smaller *τ* (faster network dynamics) because the particles are confined to linear or looped fragments for shorter time periods.

To identify different particle spread regimes we show in Fig. 2E,F the effective diffusivity and MFPT as the ratio of fusion (node connection) (fusion) and fission (disconnection) rates *λ*_add_*/λ*_remove_ is varied. For sufficiently low *λ*_add_*/λ*_remove_ the effective diffusivity (Fig. 2E) for all networks is proportional to *λ*_add_*/λ*_remove_ because particle spread is limited by the probability that a particle can cross a newly-formed fusion connection before the connection is removed (connection lifetime = 1*/λ*_remove_), as well as the rate of connection formation (∼ *λ*_add_). Similarly, for sufficiently low *λ*_add_*/λ*_remove_ the MFPT (Fig. 2F) decreases proportional to *λ*_remove_*/λ*_add_.

If *λ*_remove_ ≃ 1 for *λ*_add_*/λ*_remove_ *<* 1 then *D*_eff_ falls below and MFPT increases above the low *λ*_add_*/λ*_remove_ trend at *λ*_remove_ ≃ 1 (Fig. 2E,F). This is because particle motion across a connection is no longer limited by competition with connection lifetime (*λ*_remove_ ≲ 1) but the network remains relatively disconnected (*λ*_add_*/λ*_remove_ *<* 1). Particle spread thus falls below the low *λ*_add_*/λ*_remove_ trend, slowly increasing with the connectivity until the network becomes much more connected for *λ*_add_*/λ*_remove_ ≃ 1. At *λ*_add_*/λ*_remove_ ≃ 1 on degree-four networks the effective diffusivity rises to its maximum and MFPT falls to its minimum, remaining unchanged for further increased *λ*_add_*/λ*_remove_. If *λ*_remove_ ≃ 1 at *λ*_add_*/λ*_remove_ *>* 1, then the transition from the low *λ*_add_*/λ*_remove_ trend is directly to the maximum *D*_eff_ and minimum MFPT (Fig. 2E,F). On degree-two networks with fast dynamics (high *λ*_add_) *D*_eff_ increases to a maximum for *λ*_add_*/λ*_remove_ ≃ 1 as the network becomes more connected, before *D*_eff_ decreases at higher *λ*_add_*/λ*_remove_ as the network becomes maximally connected and freezes into linear or looped fragments (Fig. 2E, F). On degree-two networks with slow dynamics (low *λ*_add_) *D*_eff_ does not have an intermediate maximum at *λ*_add_*/λ*_remove_ ≃ 1, instead increasing to a maximum *D*_eff_. Similarly, on degree-two networks with fast dynamics MFPT decreases to a minimum for *λ*_add_*/λ*_remove_ ≃ 1 and MFPT increases for higher *λ*_add_*/λ*_remove_ as the network freezes into linear or looped fragments.

Particles on degree-four dynamic lattice networks are found to monotonically increase their effective diffusivity and decrease their search time as connectivity increases. In contrast, networks limited to a maximum node degree of two can see particles maximize effective diffusivity and minimize search time at intermediate connectivity, as the highest connectivity levels freeze linear and looped structures that decrease diffusivity and increase search time. The highest effective diffusivity and lowest search times on networks with a maximum node degree of two, which nearly match the maximum effective diffusivity and minimum search times on degree-four networks, are achieved for fast network rearrangement. Stated another way, while the maximum effective diffusivity and minimum search time on degree-four networks only modestly out-perform those on networks limiting node degree to two, the degree-four networks can achieve this high performance with substantially slower network rearrangement dynamics.

These simulations vary network dynamics timescales relative to the fixed particle motion timescale. A high *τ* can represent any scenario with particle diffusion that is faster than network dynamics, such as a scenario with a rapidly-diffusing ion in the mitochondrial matrix. A low *τ* can represent any scenario with diffusion that is much slower than network dynamics, such as a slowly-diffusing membrane-embedded complex [48].

### B Aspatial agent-based model

We now apply an existing aspatial mitochondrial network model [8] (see Fig. 1 right panel) in which each minimal mitochondrial fragment does not have a spatial position and is represented by two permanently joined nodes. Two degree-one nodes can undergo end-to-end fusion with rate *a*_1_ to form a connected degree-two node, lengthening a mitochondrial tube; a free node can undergo end-to-side fusion with a degree-two node with rate *a*_2_ to form a degree-three node, branching a mitochondrial tube; and the reverse fission processes (at rates *b*_1_ and *b*_2_ = (3*/*2)*b*_1_, respectively) also occur. Particles can move from one minimal mitochondrial fragment to attached mitochondrial fragments with a rate of *k*_m_ = 1, setting the timescale. Further simulation details are provided in the Appendix.

Figure 3A shows the mean first-passage time (MFPT) for a particle initially on a randomly-selected minimal mitochondrial fragment to find another randomly-selected target fragment as the ratio of end-to-end connections to fissions, *a*_1_*N*_mito_*/b*_1_ is varied. This ratio is analogous to the *λ*_add_*/λ*_remove_ varied in Fig. 2E,F for lattice networks. For both degree-three model mitochondrial networks that permit branching (*a*_2_ ≠ 0, solid curves in Fig. 3A) and degree-two networks that do not permit branching (*a*_2_ = 0, dashed curves in Fig. 3A), for highly fragmented networks with low *a*_1_*N*_mito_*/b*_1_ the MFPT is proportional to *b*_1_*/*(*a*_1_*N*_mito_). This trend is due to particle motion that is limited by the probability that a particle in one mitochondria crosses a newly-formed but short-lived fusion connection to another mitochondria, similar to the MFPT ∼ *λ*_remove_*/λ*_add_ trend in Fig. 2F.

For branching mitochondrial networks (solid curves in Fig. 3A), the MFPT stops decreasing along the ∼ *b*_1_*/*(*a*_1_*N*_mito_) trend for all end-to-end fusion *a*_1_ rates at *a*_1_*N*_mito_*/b*_1_ ≃ 0.1 – 1. This change in the MFPT trend is because the search time on the decreasing trend is approaching the minimum search time.

The minimum search time corresponds to the scenario that a particle will diffuse to a randomly selected mitochondrial fragment as quickly as the particle can diffuse from one mitochondrial fragment to another. The mean time to find a target mitochondrial fragment is the product of the time per attempt (diffusion to another mitochondrial fragment) and the mean number of attempts necessary to find the target fragment, ⟨*T*_min_⟩ = *τ*_attempt_⟨*N*_attempts_⟩, with *τ*_attempt_ = 1*/k*_m_. With the probability 1*/N*_mito_ that each successive attempt is successful and the probability [(*N*_mito_ − 1)*/N*_mito_]^*n−*1^ that the first *n* − 1 attempts are unsuccessful, the probability that the target fragment will be found on attempt *n* is

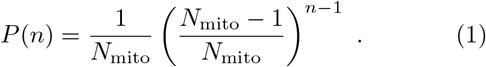

With the mean number of attempts 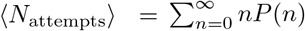 then

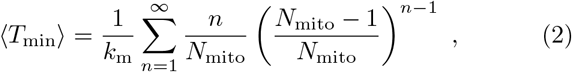

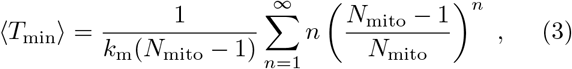

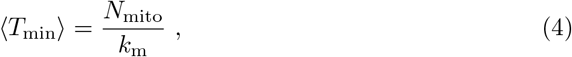

with the final line applying 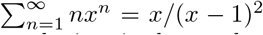 for *x <* 1. Thus the minimum search time is the product of the number of mitochondrial fragments and the timescale for particle diffusion between fragments.

For *a*_1_*N*_mito_*/b*_1_ ≳ 1 the MFPT for branching networks (solid curves in Fig. 3A) is mostly independent of *a*_1_*N*_mito_*/b*_1_, with approximately flat MFPT in Fig. 3A, because in this regime the networks are well connected with frequent end-to-end and/or many end-to-side connections, facilitating rapid search. Solid curves in Fig. 3C show the number of degree-two nodes for branching networks rises above zero at *a*_1_*N*_mito_*/b*_1_ ≃ 10^*−*2^ – 10^*−*1^. For rapid end-to-end fusion (high *a*_1_) the number of degree-two nodes continues to rise as *a*_1_*N*_mito_*/b*_1_ further increases, while for slower fusion (low *a*_1_) the number of degree-two nodes only rises somewhat before decreasing.

Degree-two nodes can undergo end-to-side fusion with a free end (degree-one node) to become a branch in the mitochondrial network (degree-three node). As shown in Fig. 3B, the value of *a*_1_*N*_mito_*/b*_1_ above which degree-three nodes increase depends on the end-to-end fusion rate *a*_1_, with degree-three nodes increasing at the *a*_1_*N*_mito_*/b*_1_ value at which degree-two nodes (solid curves) decrease in Fig. 3C, indicating that the increase in degree-two nodes is converted to an increase degree-three nodes. The *a*_1_*N*_mito_*/b*_1_ value for a branched network at which degree-two nodes decrease and degree-three nodes increase depends on the end-to-end fusion rate *a*_1_ because fusion of a degree-two node and a free end to form a degree-three node competes with fission into two free ends. The initial rise in degree-two nodes for a branched network at *a*_1_*N*_mito_*/b*_1_ ≃ 10^*−*2^ – 10^*−*1^ is because this combination represents the balance between end-to-end fusion and fission. However, the balance between a degree-two node either fusing with a free end to form a degree-three node at rate *a*_2_ or undergoing fission to form two free ends at rate *b*_1_ is controlled by the ratio of *a*_2_*/b*_1_ — at a given *a*_1_*N*_mito_*/b*_1_ in Figs. 3B,C a higher end-to-end fusion rate *a*_1_ defines a higher *b*_1_, such that increasing the number of degree-three nodes requires more degree-two nodes, which occurs at higher *a*_1_*N*_mito_*/b*_1_. We see corresponding changes in the number of free ends (degree-one nodes) in solid curves of Fig. 3D, as the number of free ends decreases when degree-two nodes increase in Fig. 3C (solid curves). Recent modeling work shows similar trends in the number of degree-three, degree-two, and degree-one nodes as the ratio of fusion to fission is varied [19].

For non-branching networks (dashed curves in Fig. 3A), the MFPT stops decreasing along the ∼ *b*_1_*/*(*a*_1_*N*_mito_) trend when *b*_1_ ≃ 1. For low *a*_1_ ≲ 0.1 s^*−*1^, as *a*_1_*N*_mito_*/b*_1_ increases above the value at which *b*_1_ ≃ 1, the MFPT is controlled by the timescale of mitochondrial dynamics, with particles waiting for new connections to search for the target mitochondrial fragment, and increasing *a*_1_ leads to lower MFPT. For higher *a*_1_ ≳ 0.1 s^*−*1^, as *a*_1_*N*_mito_*/b*_1_ increases above *b*_1_ ≃ 1, the MFPT is controlled by the timescale of particle motion, with many connections allowing the particle to continuously diffuse to another mitochondrial fragment as it undergoes a search for the target mitochondrial fragment. This corresponds to the minimum search time ⟨*T*_min_⟩ of Eq. 4. At sufficiently high *a*_1_*N*_mito_*/b*_1_ ≳ 10^2^, the MFPT for non-branching networks increases (dashed curves in Fig. 3A). This is distinct behavior in comparison to branching networks, which have a flat MFPT independent of fission and fusion rates for high *a*_1_*N*_mito_*/b*_1_.

This increasing MFPT for non-branching networks is because the network freezes into linear and looped fragments as the fission rate *b*_1_ becomes low, and the number of free ends (degree-one nodes) decreases to nearly zero (dashed curves in Fig. 3D) and nearly all nodes are part of end-to-end fusions (degree-two nodes, dashed curves in Fig. 3C) for *a*_1_*N*_mito_*/b*_1_ ≳ 10^2^. Although the degree-two and degree-one node populations for branched networks (solid curves in Fig. 3C and 3D, respectively) change as *a*_1_ is varied, without branching the degree-two and degree-one node populations (dashed curves in Fig. 3C and 3D, respectively) are independent of *a*_1_ and are determined by *a*_1_*N*_mito_*/b*_1_ as these non-branched networks are only balancing end-to-end fusion and fission.

Particles on both branching and non-branching model aspatial mitochondrial networks decrease their search time as the networks increase their connectivity from nearly entirely fragmented to well-connected networks. Once a branching network has become well-connected, further increases in fusion decreases in fission, or changes to the timescale of network dynamics have a limited effect on search time. In contrast, once a non-branching network has become well connected if fusion is sufficiently increased or fission sufficiently decreased, then MFPT substantially rises as the network becomes frozen into looped and linear fragments. For non-branching networks, the timescale of network dynamics affects the minimum MFPT reached, with fast dynamics achieving lower MFPT than slow dynamics. Non-branched networks with sufficiently fast dynamics are also able to achieve the minimum MFPT achieved by branched networks across timescales of dynamics. Overall, the minimum MFPT achieved by branching networks with all dynamical timescales can only be achieved by non-branching networks with fast dynamics, the non-branching networks with slow dynamics exhibiting substantially longer search times.

A single end-to-side fusion rate *a*_2_ is insufficient to explore the range of possible mitochondrial network connectivity, as the levels of free ends (degree-one nodes), end-to-end fusions (degree-two nodes), and tip-to-side fusions (degree-three nodes) depend nonlinearly on the fusion-fission ratios *a*_1_*/b*_1_ and *a*_2_*/b*_2_ [8]. In our simulations of aspatial mitochondrial networks with branching thus far, the end-to-side fusion rate *a*_2_ = 0.01 s^*−*1^ (Fig. 3). This *a*_2_ value is selected to provide an intermediate level of network branching, with similar *a*_2_ values (increased or decreased by an order of magnitude) similarly providing an MFPT ∼ *b*_1_*/*(*a*_1_*N*_mito_) trend for fragmented networks at low *a*_1_*N*_mito_*/b*_1_ and an unchanging MFPT for well-connected networks at higher *a*_1_*N*_mito_*/b*_1_, as shown in Fig. 3A (solid curves) and Fig. 6B,C. A substantially decreased *a*_2_ value provides less branching and leads towards the MFPT behavior for unbranched networks, with lower end-to-side fusion causing longer MFPT at a given *a*_1_*N*_mito_*/b*_1_ (Fig. 3A(dashed curves) and Fig. 6D). A substantially increased *a*_2_ value provides denser branching and leads to MFPTs that are shorter than the ∼ *b*_1_*/*(*a*_1_*N*_mito_) trend (Fig. 6A) in the nearly-disconnected network regime, with branching connecting short mitochondria with end-to-side fusions to other mitochondria, enabling a particle to more rapidly sample more mitochondria.

**FIG. 6.**
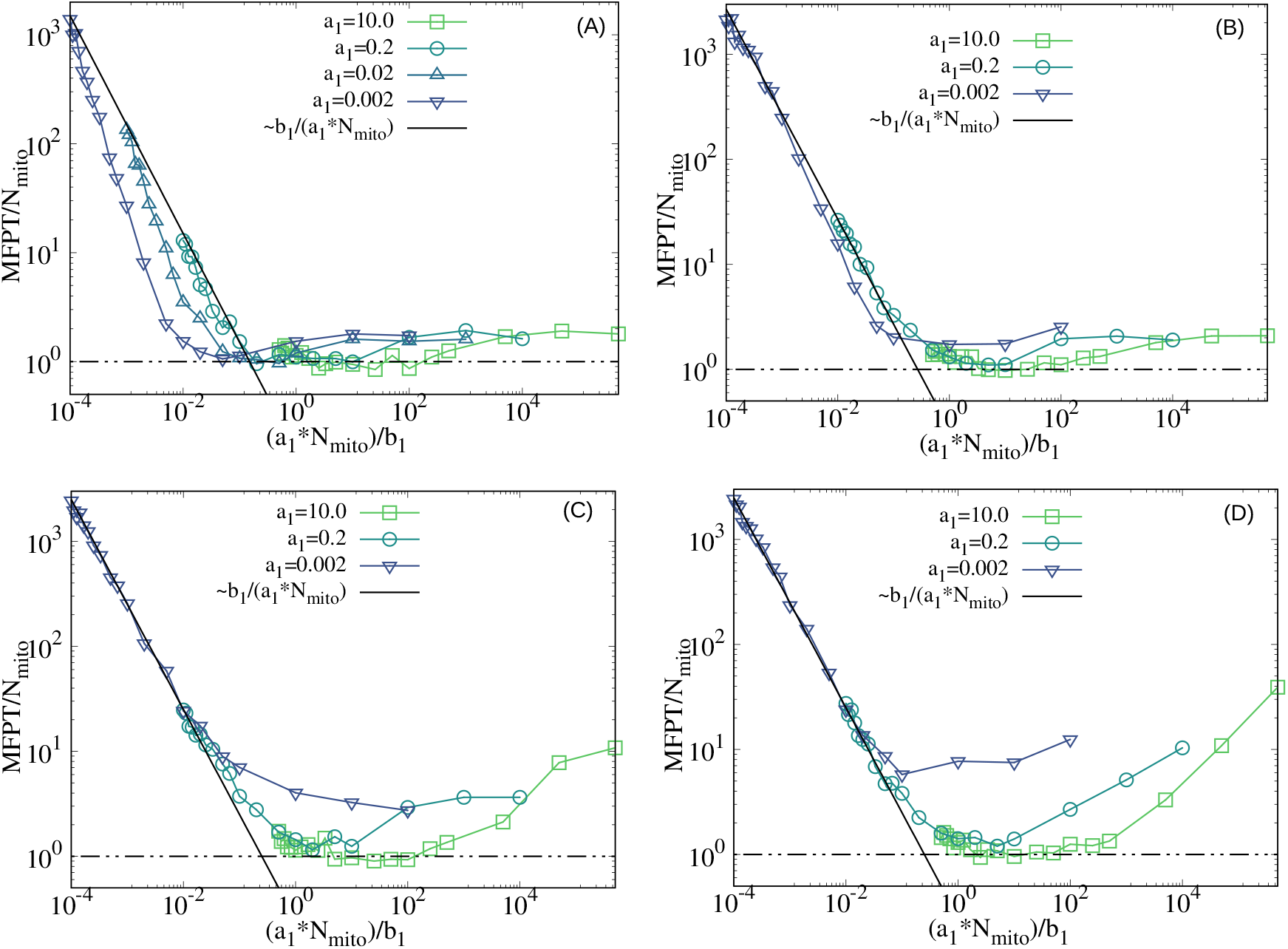
Search times for particle diffusing on model aspatial dynamic mitochondrial network with variation in end-to-side fusion levels. Mean first-passage time (MFPT) of a particle to find a randomly-selected mitochondrial fragment from an initial location on another randomly-selected fragment, as the scaled ratio of end-to-end fusion to fission *a*_1_*N*_mito_*/b*_1_ is varied. End-to-side fusion rate varies between panels with (A) *a*_2_ = 10, (B) *a*_2_ = 0.1, (C) *a*_2_ = 0.001, and (D) *a*_2_ = 0.00001. The black solid line is ∼ *b*_1_*/*(*a*_1_*N*_mito_) and the dot-dashed black line is minimum search time ⟨*T*_min_⟩ from Eq. 4. Legend indicates end-to-end fusion rate *a*_1_. Across all panels, the rate to move to connected mitochondrial fragment is *k*_m_ = 1, the number of minimal mitochondrial fragments *N*_mito_ = 50, and data averaged over 100 samples.

These simulations vary network dynamics timescales relative to the fixed rate at which particles move to connected mitochondria fragments, *k*_m_ = 1, setting the timescale. Across the *a*_1_ range explored for branching networks with *a*_2_ = 0.01 (solid curves, Fig. 3A), particle search is affected little by whether particle motion is relatively fast or slow. For the unbranched networks (dashed curves, Fig. 3A), particles with slower diffusion relative to the speed of network dynamics (large *a*_1_) lead to lower MFPT, as the mitochondrial fragment has the opportunity to rearrange connections, which could represent a protein diffusing in a mitochondrial network with rapid dynamics or a slowly diffusing protein complex in a mitochondrial network with typical dynamics. Also on unbranched networks, particles that are more diffusive relative to the speed of network dynamics (small *a*_1_) could represent a protein diffusing in a mitochondrial network with slow dynamics or a rapidly diffusing ion in mitochondria with rapid dynamics. Figure 3 explored a mitochondrial network with *N*_mito_ = 50 minimal mitochondria fragments, with larger networks (*N*_mito_ = 100) showing very similar MFPTs (Fig. 7).

**FIG. 7.**
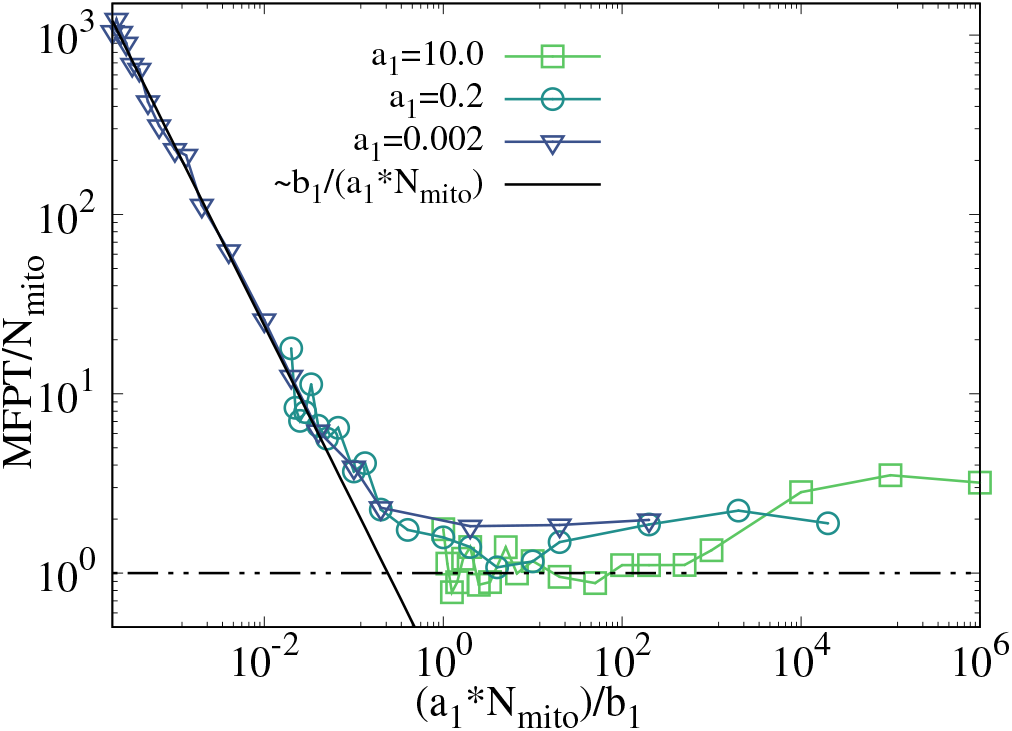
Search times for particle diffusing on model aspatial dynamic mitochondrial network with larger network size. Mean first-passage time (MFPT) of a particle to find a randomly-selected mitochondrial fragment from an initial location on another randomly-selected fragment, as the scaled ratio of end-to-end fusion to fission *a*_1_*N*_mito_*/b*_1_ is varied. Black solid line is ∼ *b*_1_*/*(*a*_1_*N*_mito_) and dot-dashed black line is minimum search time ⟨*T*_min_⟩ from Eq. 4. Legend indicates end-to-end fusion rate *a*_1_. *a*_2_ = 0.01, the rate to move to connected mitochondrial fragments is *k*_m_ = 1, number of minimal mitochondrial fragments *N*_mito_ = 100 (in contrast to *N*_mito_ = 50 for other aspatial model results), and data averaged over 100 samples.

## III DISCUSSION

Mitochondria form dynamic tubular networks that continuously remodel their structure through fusion and fission processes. These dynamics are proposed to enable the sharing of mitochondrial proteins and other molecules, evening out heterogeneous delivery of nuclear-encoded proteins and complementing mutant protein copies from nearby mitochondrial DNA copies with those from more distant copies, among other advantages from sharing molecules [31, 37, 39]. Mitochondrial networks can range from fragmented [18] to highly connected structures [9, 43], depending on the cell type and environment. We build on previous quantitative modeling of mitochondrial network dynamics, using a two-dimensional spatial lattice [31] and an aspatial agent-based model [8], to explore how fusion and fission levels control the spread of particles through the mitochondrial network.

As expected, better-connected networks and networks with faster fusion and fission dynamics each lead to increased particle spread for both spatial and aspatial networks. For branching networks, once the mitochondrial network is sufficiently well-connected further increases in connectivity do not substantially increase particle spread. This is similar to a percolation transition in two dimensions [47], matching descriptions of mitochondrial networks as approaching effectively two dimensional [9, 21, 44].

For the most fragmented networks with much faster fission than fusion, particle spread is limited by the speed of fission dynamics as fission competes with the timescale for particle movement between fused mitochondria, with particle spread increasing as fission slows. If the network becomes well connected while fission is still rapid compared to particle motion between mitochondria, then the increase in spread primarily occurs due to slowing fission competing less with but still limiting particle motion, and the network becoming well-connected halts further improvement in spread. This scenario applies to particles with relatively low diffusivity within mitochondria, such as large membrane complexes. If the fission rate instead slows to no longer compete with particle motion (particles have plenty of time to cross a recently-formed fusion connection), then on the spatial lattice-based network spread will plateau until fisson decreases sufficiently that the network becomes well-connected causing particle spread to become more rapid and reach a maximum. On the aspatial network, the particles instead never reach the fastest spread. This distinction is because on the spatial network, more connectivity always benefits spread even if network dynamics are slow, while on the aspatial network high connectivity combined with slow dynamics can lead to persistent disconnected sub-networks. This scenario, of particles easily crossing a fusion connection before fission removes the connection, applies to particles with relatively high diffusivity within mitochondria, such as ions or soluble proteins in the mitochondrial matrix.

Experimental measurements of protein spread on mitochondrial networks support these two regimes of particle spread, one regime limited by fission events occurring before particles can cross the fused connection and the other regime defined by fusion event lifetimes much longer than the timescale for particles to cross the fused connection. While fusion events with long lifetimes enable ongoing protein spread, short-lived mitochondrial ‘kiss-and-run’ fusion events also allow substantial exchange of soluble proteins but limited exchange of membrane proteins between mitochondrial fragments [49], and can enable meaningful protein delivery for extended mitochondrial networks [50]. Soluble proteins are also observed to spread through yeast mitochondrial networks in minutes following fusion, while protein components of the oxidative phosphorylation machinery, which form membrane-embedded complexes and can be largely confined to cristae, show limited spread through the network more than an hour after fusion [51]. Beyond molecule size and membrane embedding, which can influence diffusivity in mitochondria, various protein-specific and environmental variables influence diffusivity patterns [52].

An important mitochondrial network feature is their three-way junctions, formed via end-to-side fusion, as they enable branching and looping of mitochondrial networks [9–11]. Our model explores both branching model mitochondrial networks with three-way junction formation and non-branching networks that are not permitted to form three-way junctions. Particle spread on non-branching model mitochondrial networks (described above) is modified for non-branching networks. Diffusive particles on both branching and non-branching networks can achieve comparably fast spread, however, non-branching networks require much faster fusion and fission dynamics to achieve the most rapid spread. This suggests that one advantage of branching mitochondrial networks is to enable the efficient spread of proteins and other molecules with much lower costs associated with the very rapid fusion and fission that would be necessary for efficient spread in a non-branching mitochondrial network. Furthermore, spread on non-branching mitochondrial networks does not remain maximized once networks are sufficiently well-connected. Instead, non-branching networks have an optimum connectivity that maximizes spread, with both lower and higher connectivity decreasing spread. While less spread for less connectivity is similar to branching networks, more connectivity causes decreased spread on non-branching networks because the network freezes into linear fragments and loops.

The mitochondria modeled as a spatial network on a two-dimensional lattice and as an aspatial agent-based network differ in how particle spread changes as fission slows compared to fusion (described above) and in how spread on non-branching networks approaches spread on branching networks. For spatial networks, the particle spread timescale on non-branching networks does not achieve the lower spread timescale of branching networks, while for aspatial networks the spread timescale on non-branching networks (with fast dynamics and optimum connectivity) can reach the spread timescale on branching networks. This distinction, with spread on non-branched spatial networks persistently disadvantaged relative to branched networks, demonstrates the important effect of embedding a network in physical space [53]. Rapid dynamics does not entirely rescue the slow spread on non-branching spatial lattice-based mitochondrial networks because mitochondrial fragments embedded on a spatial lattice are constrained to connect to the same few nearby fragments. Very recent work developed a mitochondrial network model permitting mitochondrial fragment motion, integrating the spatial aspect of our lattice model with the changing fusion partners of our aspatial model [19]. Although that work did not examine particle spread, frequent network rearrangements were found to require intermediate fusion rates, to allow recently fissioned mitochondrial fragments to separate without refusing, suggesting the spatially embedded nature of mitochondria constrains network structure and downstream processes such as spread.

Yeast mitochondria are well-connected into a small number of network components [9], while mammalian mitochondria can be near the percolation transition threshold with a range of fragment sizes [8, 43]. We then expect that yeast are typically near optimized spread and a substantial shift towards fragmentation via more fission or less fusion would correspondingly slow spread through the mitochondrial network. Mammalian mitochondria are instead in the regime with spread affected by smaller changes to connectivity. However, mammalian mitochondria have relatively fast dynamics [19], such that for many proteins and other molecules spread may be limited by competition between intra-mitochondrial diffusion and fission, a regime that yields faster spread across parameter values. In contrast to both yeast and mammals, plant mitochondrial networks are very fragmented [23], suggesting spread through plant mitochondria is significantly hindered compared to better-connected networks, and sensitive to the frequency and lifetime of fusion connections.

Through quantitative modeling of spread on mitochondrial networks, we have identified network branching at three-way junctions as crucial to allow efficient spread with relatively slow mitochondrial fusion and fission dynamics. We have also shown that spread falls into different regimes that correspond to regimes of network connectivity: on fragmented networks spread is slow and depends on marginal changes to fusion and fission, while on well-connected networks spread is rapid and depends little on marginal changes to fusion and fission. By exploring both a spatial and aspatial model of mitochondrial dynamics, our results align with previous work suggesting that the spatial nature of mitochondria is important to network rearrangements. This work advances quantitative understanding of mitochondrial network structure and dynamics towards mechanistic connection to mitochondrial function.

## ACKNOWLEDGMENTS

This work was supported by a Natural Sciences and Engineering Research Council of Canada (NSERC) Discovery Grant, by start-up funds provided by the Toronto Metropolitan University Faculty of Science, by the Toronto Metropolitan University Faculty of Science Dean’s Research Fund, and was enabled by computational resources provided by the Digital Research Alliance of Canada (alliancecan.ca), including those provided as part of a Resource Allocation cluster usage resource allocation.

## Appendix A: Simulation details

### 1 Spatial lattice-based model

The mitochondrial network is modeled as a square 2D lattice structure, with mitochondrial nodes on lattice sites that can be linked to their nearest neighbors by edges. Fusion is represented by the addition of an edge between two nearest-neighbor nodes with rate *λ*_add_ and fission as the removal of an edge with rate *λ*_remove_. *p* = *λ*_add_*/*(*λ*_add_ + *λ*_remove_) is the probability of a particular edge to be present and *τ* = 1*/*(*λ*_add_ + *λ*_remove_) is the timescale of network dynamics. A particle at a node has a rate of one to move to each nearest neighbor node connected by an edge. Model network dynamics and particle motion are simulated by using the Gillespie algorithm [54]. For *τ* = 0 the status of each edge is not tracked, and instead, when the particle attempts to cross to another node the edge is present (and the particle allowed to cross) with probability *p*. For *τ* = ∞ each edge is initially present with probability *p*, with no further edge dynamics. This model is similar to that described previously [31].

### 2 Aspatial agent-based model

This model does not assign spatial positions to mitochondria. Each minimal mitochondrial fragment is modeled as two permanently connected nodes. A mitochondrial network is composed of three different types of nodes: free nodes (degree-one nodes), degree-two nodes, and branching nodes (degree-three nodes). Two free nodes can end-to-end fuse to form a degree-two node with rate *a*_1_ and a degree-two node can fission into two free nodes with rate *b*_1_. A degree-two node can end-to-side fuse to a free node to form a degree-three node with rate *a*_2_ and a degree-three node can fission into a degree-two node and a free node with rate *b*_2_ (the node which becomes free is randomly selected). Nodes may not fuse to their permanently-connected partner node. A particle on a particular minimal mitochondrial fragment can move to any connected mitochondrial fragment with rate *k*_m_ = 1, with the specific mitochondrial fragment chosen to move to randomly selected. Model network dynamics and particle motion are simulated by using the Gillespie algorithm [54]. Most details of this model are from previous work [8].

## Appendix B: Additional results

### 1 Spatial lattice-based model

The effective diffusivity *D*_eff_ vs connection probability *p*, as in Fig. 2A but compared to a larger system is shown in Fig. 4A. The larger system behaves nearly identically to the smaller network. Figure 4B similarly compares effective diffusivity for both degree-four and degree-two networks for a larger system compared to the system for results in Fig. 2C, finding the larger system behaves nearly identically to the smaller network.

Figure 5 compares effective diffusivity *D*_eff_ and mean first-passage time (MFPT) of a particle on a dynamic spatial lattice network for the case with nodes only allowed to connect to three nearest neighbors (degree-three networks) to the case where nodes can connect to all four nearest neighbors (degree-four networks, Fig. 2A). The qualitative *D*_eff_ and MFPT are unchanged, although *D*_eff_ substantially decreases and MFPT substantially increases at high *λ*_add_*/*(*λ*_add_ +*λ*_remove_) for degree-three networks compared to degree-four networks. This regime in which these networks differ is when the dynamics favor high connectivity and thus the limit to three connections shows an effect.

### 2 Aspatial agent-based model

Figure 3A examines the mean first-passage time (MFPT) for the aspatial agent-based dynamic mitochondrial network model for a branching network with intermediate end-to-side fusion rate *a*_2_ = 0.01 and a non-branching network with *a*_2_ = 0. Figure 6 explores other *a*_2_ values. For lower *a*_2_ values (Fig. 6C,D) the MFPT trends towards the results for the non-branching model (Fig. 3A, dashed curves). For higher *a*_2_ values (Fig. 6A,B) the MFPT decreases below the MFPT ∼ *b*_1_*/*(*a*_1_*N*_mito_) trend as branching fusions connecting short mitochondrial fragments and allow a particle to sample mitochondria rapidly. Since three-way junctions fission into a free end and a degree-two node at rate *b*_2_ = (3*/*2)*b*_1_, the high *a*_2_ value allows the MFPT to decrease below the MFPT ∼ *b*_1_*/*(*a*_1_*N*_mito_) trend.

Figure 7 compares MFPT for a network with *N*_mito_ = 100 mitochondrial fragments instead of *N*_mito_ = 50 in Fig. 3A, showing very similar search times.

## References

[1] B. Westermann, Mitochondrial fusion and fission in cell life and death, Nature Reviews Molecular cell biology 11, 872 (2010).

[2] A. S. Monzel, J. A. Enríquez, and M. Picard, Multifaceted mitochondria: moving mitochondrial science beyond function and dysfunction, Nature Metabolism 5, 546 (2023).

[3] O. Bergman and D. Ben-Shachar, Mitochondrial oxidative phosphorylation system (oxphos) deficits in schizophrenia: possible interactions with cellular processes, The Canadian Journal of Psychiatry 61, 457 (2016).

[4] M. Liesa, M. Palacín, and A. Zorzano, Mitochondrial dynamics in mammalian health and disease, Physiological reviews 89, 799 (2009).

[5] J. Zhao, J. Zhang, M. Yu, Y. Xie, Y. Huang, D. W. Wolff, P. W. Abel, and Y. Tu, Mitochondrial dynamics regulates migration and invasion of breast cancer cells, Oncogene 32, 4814 (2013).

[6] N. Exner, A. K. Lutz, C. Haass, and K. F. Winklhofer, Mitochondrial dysfunction in parkinson’s disease: molecular mechanisms and pathophysiological consequences, The EMBO journal 31, 3038 (2012).

[7] E. Tönnies and E. Trushina, Oxidative stress, synaptic dysfunction, and alzheimer’s disease, Journal of Alzheimer’s Disease 57, 1105 (2017).

[8] V. M. Sukhorukov, D. Dikov, A. S. Reichert, and M. Meyer-Hermann, Emergence of the mitochondrial reticulum from fission and fusion dynamics, PLoS computational biology 8, e1002745 (2012).

[9] M. P. Viana, A. I. Brown, I. A. Mueller, C. Goul, E. F. Koslover, and S. M. Rafelski, Mitochondrial fission and fusion dynamics generate efficient, robust, and evenly distributed network topologies in budding yeast cells, Cell systems 10, 287 (2020).

[10] A. I. Brown, L. M. Westrate, and E. F. Koslover, Impact of global structure on diffusive exploration of organelle networks, Scientific reports 10, 4984 (2020).

[11] G. R. Lewis and W. F. Marshall, Mitochondrial networks through the lens of mathematics, Physical biology 20, 051001 (2023).

[12] T. Kleele, T. Rey, J. Winter, S. Zaganelli, D. Mahecic, H. Perreten Lambert, F. P. Ruberto, M. Nemir, T. Wai, T. Pedrazzini, et al., Distinct fission signatures predict mitochondrial degradation or biogenesis, Nature 593, 435 (2021).

[13] M. Liesa and O. S. Shirihai, Mitochondrial dynamics in the regulation of nutrient utilization and energy expenditure, Cell metabolism 17, 491 (2013).

[14] S. Campello, R. A. Lacalle, M. Bettella, S. Mañes, L. Scorrano, and A. Viola, Orchestration of lymphocyte chemotaxis by mitochondrial dynamics, The Journal of experimental medicine 203, 2879 (2006).

[15] B. J. Seo, S. H. Yoon, and J. T. Do, Mitochondrial dynamics in stem cells and differentiation, International journal of molecular sciences 19, 3893 (2018).

[16] S. Hoppins, L. Lackner, and J. Nunnari, The machines that divide and fuse mitochondria, Annu. Rev. Biochem. 76, 751 (2007).

[17] R. E. Jensen, A. E. Aiken Hobbs, K. L. Cerveny, and H. Sesaki, Yeast mitochondrial dynamics: fusion, division, segregation, and shape, Microscopy research and technique 51, 573 (2000).

[18] Z. Wang, P. Natekar, C. Tea, S. Tamir, H. Hakozaki, and J. Schöneberg, Mitotnt: Mitochondrial temporal network tracking for 4d live-cell fluorescence microscopy data, PLOS Computational Biology 19, e1011060 (2023).

[19] K. Holt, J. Winter, S. Manley, and E. F. Koslover, Spatiotemporal modeling of mitochondrial network architecture, bioRxiv 10.1101/2024.01.24.577101 (2024).

[20] A. R. Fenton, T. A. Jongens, and E. L. Holzbaur, Mitochondrial dynamics: Shaping and remodeling an organelle network, Current opinion in cell biology 68, 28 (2021).

[21] P. K. Patel, O. Shirihai, and K. C. Huang, Optimal dynamics for quality control in spatially distributed mitochondrial networks, PLoS Computational Biology 9, e1003108 (2013).

[22] M. Guo, A. J. Ehrlicher, M. H. Jensen, M. Renz, J. R. Moore, R. D. Goldman, J. Lippincott-Schwartz, F. C. Mackintosh, and D. A. Weitz, Probing the stochastic, motor-driven properties of the cytoplasm using force spectrum microscopy, Cell 158, 822 (2014).

[23] J. M. Chustecki, D. J. Gibbs, G. W. Bassel, and I. G. Johnston, Network analysis of arabidopsis mitochondrial dynamics reveals a resolved tradeoff between physical distribution and social connectivity, Cell systems 12, 419 (2021).

[24] R. Rossignol, R. Gilkerson, R. Aggeler, K. Yamagata, S. J. Remington, and R. A. Capaldi, Energy substrate modulates mitochondrial structure and oxidative capacity in cancer cells, Cancer research 64, 985 (2004).

[25] A. J. Molina, J. D. Wikstrom, L. Stiles, G. Las, H. Mohamed, A. Elorza, G. Walzer, G. Twig, S. Katz, B. E. Corkey, et al., Mitochondrial networking protects β-cells from nutrient-induced apoptosis, Diabetes 58, 2303 (2009).

[26] L. C. Gomes, G. D. Benedetto, and L. Scorrano, During autophagy mitochondria elongate, are spared from degradation and sustain cell viability, Nature cell biology 13, 589 (2011).

[27] S. M. Rafelski, Mitochondrial network morphology: building an integrative, geometrical view, BMC biology 11, 1 (2013).

[28] H. Zhang, K. J. Menzies, and J. Auwerx, The role of mitochondria in stem cell fate and aging, Development 145, dev143420 (2018).

[29] T. Tsuboi, M. P. Viana, F. Xu, J. Yu, R. Chanchani, X. G. Arceo, E. Tutucci, J. Choi, Y. S. Chen, R. H. Singer, et al., Mitochondrial volume fraction and translation duration impact mitochondrial mrna localization and protein synthesis, Elife 9, e57814 (2020).

[30] G. López-Doménech and J. T. Kittler, Mitochondrial regulation of local supply of energy in neurons, Current Opinion in Neurobiology 81, 102747 (2023).

[31] H. Hoitzing, I. G. Johnston, and N. S. Jones, What is the function of mitochondrial networks? a theoretical assessment of hypotheses and proposal for future research, Bioessays 37, 687 (2015).

[32] J. M. Chustecki and I. G. Johnston, Collective mitochondrial dynamics resolve conflicting cellular tensions: From plants to general principles, in Seminars in Cell & Developmental Biology, Vol. 156 (Elsevier, 2024) pp. 253–265.

[33] I. Scott and R. J. Youle, Mitochondrial fission and fusion, Essays in biochemistry 47, 85 (2010).

[34] C.-R. Chang and C. Blackstone, Dynamic regulation of mitochondrial fission through modification of the dynamin-related protein drp1, Annals of the new York Academy of Sciences 1201, 34 (2010).

[35] C. C. Williams, C. H. Jan, and J. S. Weissman, Targeting and plasticity of mitochondrial proteins revealed by proximity-specific ribosome profiling, Science 346, 748 (2014).

[36] X. G. Arceo, E. F. Koslover, B. M. Zid, and A. I. Brown, Mitochondrial mrna localization is governed by translation kinetics and spatial transport, PLoS computational biology 18, e1010413 (2022).

[37] A. H. Khan, R. J. Patel, M. P. Viana, S. M. Rafelski, A. I. Brown, B. M. Zid, and T. Tsuboi, Mitochondrial protein heterogeneity stems from the stochastic nature of co-translational protein targeting in cell senescence, bioRxiv, 2023 (2023).

[38] J. M. Palozzi, S. P. Jeedigunta, and T. R. Hurd, Mitochondrial dna purifying selection in mammals and invertebrates, Journal of molecular biology 430, 4834 (2018).

[39] T. Ono, K. Isobe, K. Nakada, and J.-I. Hayashi, Human cells are protected from mitochondrial dysfunction by complementation of dna products in fused mitochondria, Nature genetics 28, 272 (2001).

[40] M. P. Viana, S. Lim, and S. M. Rafelski, Quantifying mitochondrial content in living cells, in Methods in cell biology, Vol. 125 (Elsevier, 2015) pp. 77–93.

[41] F. M. Fazal, S. Han, K. R. Parker, P. Kaewsapsak, J. Xu, A. N. Boettiger, H. Y. Chang, and A. Y. Ting, Atlas of subcellular rna localization revealed by apex-seq, Cell 178, 473 (2019).

[42] H. S. Ilamathi, M. Ouellet, R. Sabouny, J. Desrochers-Goyette, M. A. Lines, G. Pfeffer, T. E. Shutt, and M. Germain, A new automated tool to quantify nucleoid distribution within mitochondrial networks, Scientific reports 11, 22755 (2021).

[43] N. Zamponi, E. Zamponi, S. A. Cannas, O. V. Billoni, P. R. Helguera, and D. R. Chialvo, Mitochondrial network complexity emerges from fission/fusion dynamics, Scientific reports 8, 1 (2018).

[44] N. Zamponi, E. Zamponi, S. A. Cannas, and D. R. Chialvo, Universal dynamics of mitochondrial networks: a finite-size scaling analysis, Scientific Reports 12, 17074 (2022).

[45] V. M. Sukhorukov and M. Meyer-Hermann, Structural heterogeneity of mitochondria induced by the microtubule cytoskeleton, Scientific reports 5, 13924 (2015).

[46] K. Giannakis, J. M. Chustecki, and I. G. Johnston, Exchange on dynamic encounter networks allows plant mitochondria to collect complete sets of mitochondrial dna products despite their incomplete genomes, Quantitative Plant Biology 3, e18 (2022).

[47] D. Stauffer and A. Aharony, Introduction to percolation theory(CRC press, 2018).

[48] V. Wilkens, W. Kohl, and K. Busch, Restricted diffusion of oxphos complexes in dynamic mitochondria delays their exchange between cristae and engenders a transitory mosaic distribution, Journal of cell science 126, 103 (2013).

[49] X. Liu, D. Weaver, O. Shirihai, and G. Hajnóczky, Mitochondrial ‘kiss-and-run’: interplay between mitochondrial motility and fusion–fission dynamics, The EMBO journal 28, 3074 (2009).

[50] A. Agrawal and E. F. Koslover, Optimizing mitochondrial maintenance in extended neuronal projections, PLOS Computational Biology 17, e1009073 (2021).

[51] C. Jakubke, R. Roussou, A. Maiser, C. Schug, F. Thoma, D. Bunk, D. Hörl, H. Leonhardt, P. Walter, T. Klecker, et al., Cristae-dependent quality control of the mitochondrial genome, Science advances 7, eabi8886 (2021).

[52] V. M. Sukhorukov, D. Dikov, K. Busch, V. Strecker, I. Wittig, and J. Bereiter-Hahn, Determination of protein mobility in mitochondrial membranes of living cells, Biochimica et Biophysica Acta (BBA)-Biomembranes 1798, 2022 (2010).

[53] M. Pósfai, B. Szegedy, I. Bačić, L. Blagojević, M. Abért, J. Kertész, L. Lovász, and A.-L. Barabási, Impact of physicality on network structure, Nature Physics 20, 142 (2024).

[54] D. T. Gillespie, Exact stochastic simulation of coupled chemical reactions, The journal of physical chemistry 81, 2340 (1977).

